# Ultrastructural identification and molecular characterization of two new parabasalid species that naturally colonize laboratory mice, *Tritrichomonas musculus* and *Tritrichomonas casperi*

**DOI:** 10.1101/2023.01.20.524969

**Authors:** Leila Tuzlak, Eliza V.C. Alves-Ferreira, Andrea Kennard, Christina Shehata, Cindi L. Schwartz, Michael E. Grigg

## Abstract

*Tritrichomonas muris* is a flagellated protist isolated from the cecum of wild mice in the Czech Republic. This commensal protist has been shown previously to alter immune phenotypes in laboratory mice. Other trichomonads, previously referred to as *Tritrichomonas musculis* and *Tritrichomonas rainier*, also naturally colonize laboratory mice and cause immune alterations. This report formally describes two new trichomonads, *Tritrichomonas musculus* n. sp., and *Tritrichomonas casperi* n. sp., at the ultrastructural and molecular level. These two protists were isolated from laboratory mice, and were differentiated by their size and the structure of their undulating membrane and posterior flagellum. Analysis at the *18S rRNA* and trans-*ITS* genetic loci supported their designation as distinct species, related to *T. muris*. To further assess the true extent of parabasalid diversity infecting laboratory mice, 135 mice were screened at the NIH using pan-parabasalid primers that amplify the trans-*ITS* region. Forty-four percent of mice were positive for parabasalids, encompassing a total of 8 distinct sequence types. *Tritrichomonas casperi* and *Trichomitus-*like protists were dominant. *T. musculus* and *T. rainier* were also detected, but *T. muris* was not. Our work establishes a previously underappreciated diversity of commensal trichomonad protists that naturally colonize the enteric cavity of laboratory mice.

## Introduction

The gastrointestinal tract of warm-blooded vertebrates is host to a diverse microbiome composed of bacterial, viral, fungal, helminth and protist communities. Emerging evidence suggests that the microeukaryotic component of the microbiome, referred to as the eukaryome (Lukes et al. 2015), plays an important role shaping both bacterial community structure and mucosal immune homeostasis. Indeed, microbiome studies involving a number of enteric protists suggest that mutualistic protist-bacterial relationships are common and that protists actively alter microbial community structure. In healthy individuals colonized by protists within the genus *Blastocystis*, a more diverse microbiome has been described, with only some subtypes inducing gastrointestinal inflammation (Nieves-Ramirez et al. 2018). But in other cases, such as bacterial-*Entamoeba histolytica* interactions, a profound systemic disease can occur, due to the degradation of the host’s intestinal mucosal barrier (Bar et al. 2015). Other flagellated protists, including *Enteromonas, Retortamonas, Dientamoeba, Pentatrichomonas* and *Chilomastix* are generally considered enteric commensals, but some, including the parabasalid *Tritrichomonas muris* and the diplomonad *Giardia intestinalis* have been implicated in causing immune alterations in laboratory mice (Howitt et al. 2016; Solaymani-Mohammadi 2022). Protists overall appear to play important ecological roles within their hosts, highlighting a practical need to better understand the role of the eukaryome’s commensal compartment (Lukes et al. 2015).

Trichomonads belong to a group of anaerobic protists known as parabasalids, which are recognized for the unique morphology of their flagella and for having hydrogenosomes in place of mitochondria. Parabasalids are generally classified into two orders, a much larger, multiflagellated Hypermastigida that colonize wood-eating termites and cockroaches that form mutualistic protist-host relationships in the gut of their insect hosts (Ceza et al. 2015), and a smaller (<20 micron) Trichomonadida that possess six or fewer flagella and are typically commensals that infect a diverse array of vertebrates. Parabasalid species are further divided into seven sub-groups, referred to as the Cristamonadida, Honibergiellida, Hypotrichomonadida, Spirotrichonymphida, Trichomonadida, Trichonymphida, and Tritrichomonadida (Cepicka et al. 2010). The parabasalids that colonize the enteric cavity of warm-blooded vertebrates are less well studied and the true extent of their genetic diversity is unknown.

*Tritrichomonas muris* is considered the primary flagellated parabasalid that naturally colonizes mice. Its presence can drive proinflammatory T helper 1 (Th1) T cell responses in the murine gut that can exacerbate disease in colitis models (Escalante et al. 2016) as well as induce strong immunomodulating T helper 2 (Th2) T cell responses capable of altering mucosal immune homeostasis (Howitt et al. 2016). Two other sequence types have been described that naturally colonize laboratory mice, these have been referred to as *Tritrichomonas musculis* and *Tritrichomonas rainier*, but neither have been rigorously characterised at the ultrastructural or molecular level, so it remains unclear if they are separate species, distinct from *T. muris*. Whether host immunity against these various trichomonad sequence types is protist specific is not well studied. Both *T. musculis* and *T. muris* are associated with increases in ILC2 cells producing IL-5 or IL-13 in the colon (Chudnovskiy et al. 2016; Howitt et al. 2016), but *T. muris* is not associated with an upregulation of IL-17 or IL-18, whereas *T. musculis* is (Escalante et al. 2016; Chudnovskiy et al. 2016). However, both parasites have been shown to activate the inflammasome, protect mice against *Salmonella typhimurium* infection, and secrete succinate which causes intestinal tuft cell expansion (Chudnovskiy et al. 2016; Nadjsombati et al. 2018). This study was pursued to determine the true extent of genetic diversity and number of distinct trichomonad species that naturally colonize the enteric cavity of laboratory rodents.

Trichomonad blooms in the guts of laboratory mice are commonly observed by microscopy, but no standardized approach to limit their transmission or expansion in mouse litters is performed because they are considered commensal organisms (Treuting et al. 2012). Our study herein identified a spectrum of distinct parabasalids that varied in size, morphology, and their flagella number and arrangement in the ceca of laboratory mice. Purification of two of these flagellates from mono-colonized mice by flow cytometry, followed by ultrastructural and molecular characterization, confirmed that a FACS-purified protist measuring ~10 microns in length that we refer to as *T. musculus n. sp*. was 100% identical to the sequence originally described by Chudnovskiy et al (2016) as *T. musculis*, whereas a FACS-purified protist measuring ~5 microns in length, that we refer to as *T. casperi n. sp*., was 100% identical to a sequence type submitted to GenBank and isolated from C57BL/6 mice in China (Genbank Acc. No. MF375342). Combining ultrastructural data with the molecular analyses supported the designation of these two trichomonads as new species, distinct from *T. muris*. Screening laboratory mice at the NIH established that *T. casperi* was the dominant trichomonad detected, *T. musculus* and *T. rainier* were also present in the dataset. Four additional sequence types were detected that were distinct from the *Tritrichomonas* genus and require further characterization. *Tritrichomonas muris* was not detected in any of the mice. Our study herein confirms that a diverse community of commensal protists exist in the NIH laboratory mice.

## MATERIALS AND METHODS

### Purification of *T. musculus* and *T. casperi* from Cecal Content

Cecal contents of a single mono-colonized laboratory mouse bearing a single protist were harvested at 3 weeks post-gavage, soaked in a 40 mL sterile solution of 1x PBS followed by filtering through a 70 μm cell strainer. The suspension was centrifuged at 450 x*g* for 5 min and the supernatant poured off. The pellet underwent two more washes with 40 mL of PBS. Following the washes, a percoll gradient was performed to isolate the protozoa. Briefly, in a 15 mL conical centrifuge tube, the pellet was resuspended in 6 mL of 40% percoll diluted in PBS, followed by an underlay of 4 mL of 80% percoll diluted in PBS. The gradient was centrifuged with the brake off at 1000 x*g* for 15 min, after which the trichomonads were isolated by pipetting from the middle layer and transferred into a 15 mL conical. The percoll was removed by adding 12 mL PBS and then the suspension was centrifuged at 450 x*g* for 5 min and the supernatant poured off. The pellet was resuspended in 1 mL of PBS and sorted using a FACS Diva 8.0.1.

### Scanning electron microscopy

Sorted trichomonads or cecum from mono-colonized mice were prepared for scanning electron microscopy (SEM) via a Biowave microwave-assisted (Ted Pella, Inc.) protocol as described in (Jackson et al. 2018). Briefly, samples were fixed in 2% paraformaldehyde, 2.5% glutaraldehyde, 0.05% alcian blue in 0.1 M Sorenson’s phosphate buffer (Karnovsky’s Fixative, Electron Microscopy Sciences) and stored at 4 °C until further processed. Sorted cells were adhered to poly-L-lysine coated silicon wafer chips for 30 minutes. Both sorted cells and cecum pieces were post-fixed in 0.5% OsO_4_, 0.8% K_4_Fe(CN)_6_ in 0.1 M sodium cacodylate buffer for 1 hour. The samples were rinsed in dH_2_0, stained with 1% aqueous tannic acid for 1 hour, rinsed with dH_2_0, and finally fixed with a second round of 0.5% OsO_4_, 0.8% K_4_Fe(CN)_6_ in 0.1 M sodium cacodylate buffer for 1 hour to enhance conductivity under the scanning electron beam. Samples were rinsed with dH_2_0, then dehydrated using a graduated ethanol series into 100% ethanol, critical point dried using a Bal-Tec CPD 030 (Leica Microsystems, Buffalo Grove, IL), placed on a stub using carbon sticky tape, and sputter coated with 10Å of iridium (EMS300TD, Electron Microscopy Sciences). Samples were placed in a Hitachi SU8000 scanning electron microscope operating at 5.0 kV, 10 μA, and a working distance of ~8 mm using a secondary electron detector.

### DNA extraction, PCR amplification, and Sanger sequencing

DNA was isolated from sorted protists using the QIAGEN DNeasy Blood and Tissue Kit, whereas DNA was isolated from fecal samples using the QIAGEN DNeasy PowerSoil Pro Kit. PCR amplification of the *18S rRNA* gene sequence was performed using primers PF1 [tgcgctacctggttgatcctgcc] and FAD4 [tgatccttctgcaggttcacctac] (Keeling 2002) on extracted DNA. PCR amplification of the trans-*ITS* gene sequence was performed using pan-parabasalid primers in a nested reaction: 18S EXT FWD [aatacgtccctgccctttgt] and 28S EXT REV [tcctccgcttaatgagatgc] followed by 18S INT FWD [aacctgccgttggatcagt] and 28S INT REV [cttcagttcagcgggtcttc] (Chudnovskiy et al. 2016). All PCR reactions consisted of 35 cycles accordingly: 95°C for 40 seconds, 58°C for 40 seconds, 72°C for 40 seconds. Prior to sequencing, amplification of the nested PCR product was confirmed by 0.8% agarose gel electrophoresis, and confirmed positives were prepared for Sanger sequencing of the PCR population using ExoSAP-IT. Sequencing was performed by the Rocky Mountain Laboratory Genomics Unit DNA Sequencing Center, NIAID, NIH, Hamilton, Montana. Forward and reverse reads were DeNovo assembled and analyzed by BLAST using default settings in Geneious prime version 2022.0.2. Mixed infections were identified by the presence of di-nucleotides present in the DNA sequence electropherograms at specific positions known to vary between the 8 sequence types characterized. To confirm the presence of a mixed infection, the caecum from mice suspected to be co-infected with two different protists by PCR-DNA-Seq were isolated, and examined for the presence of two distinct protists morphologically, by flow cytometry, and by DNA extraction of the flow-sorted protists to repeat the PCR DNA-Seq analysis to confirm the results.

### Phylogenetic Analyses

Trichomonad sequences were aligned using the MUSCLE alignment tool (Edgar 2004), followed by the addition of other parabasalid genera to the alignment using Clustal Omega (Sievers and Higgins 2018; Sievers et al. 2011; Sievers F 2020) The alignment was manually edited, and overhangs trimmed prior to tree construction. Trees were generated in MEGA X (Stecher, Tamura, and Kumar 2020) via the Maximum Likelihood statistical method, 1000 bootstrap replications, using the Tamura-Nei model.

## RESULTS and DISCUSSION

### Morphology and Phylogeny of *T. musculus* n. sp., and *T. casperi* n. sp

Two protists of different sizes were FACS-purified from the cecal contents of laboratory mice bred in the NIH animal facility. The larger protist (referred to as *T. musculus*) was about 10 μm in length, whereas the smaller protist (referred to as *T. casperi*) was 5μm long (Fig. 1). The morphology of the undulating membrane and posterior flagellum was distinct between the two protists (Fig. 1). The undulating membrane of *T. musculus* spiraled around the length of the protist’s body (Fig. 1A, B). At the posterior end, the undulating membrane wrapped around the axostyle which supported the posterior flagellum (Fig. 1A, B). In contrast, the undulating membrane of *T. casperi* ran along the dorsal flank of the protist in a more linear fashion than observed in *T. musculus* (Fig. 1D, E). The posterior end of the undulating membrane detached from the surface and continued posteriorly as a free flagellum, which pivoted away from the main ovaloid body and axostyle (Fig. 1D, E). *In situ*, both *T. musculus* and *T. casperi* were found in giant clumps consisting of bacteria/protist aggregates at the surface of the large villi in the cecum of colonized mice (Fig. 1C, F). In liquid medium, *T. musculus* moved in a smooth, gliding fashion with the undulating membrane beating in a rhythmic manner. In contrast, *T. casperi* moved in a stochastic, apparently erratic fashion that was quite distinct from *T. musculus*. Both trichomonads differed from *T. muris* in both size and movement. *T. muris* is much larger, measuring anywhere from 16-26 microns in length, and 10-14 microns in width, and moves in a wobbly manner, distinct from the smooth gliding pattern characteristic of *T. musculus*. In similar, however, *T. muris* and *T. musculus* share a stiff, rod-like axyostyle which protrudes terminally like a tail.

**FIGURE 1.**
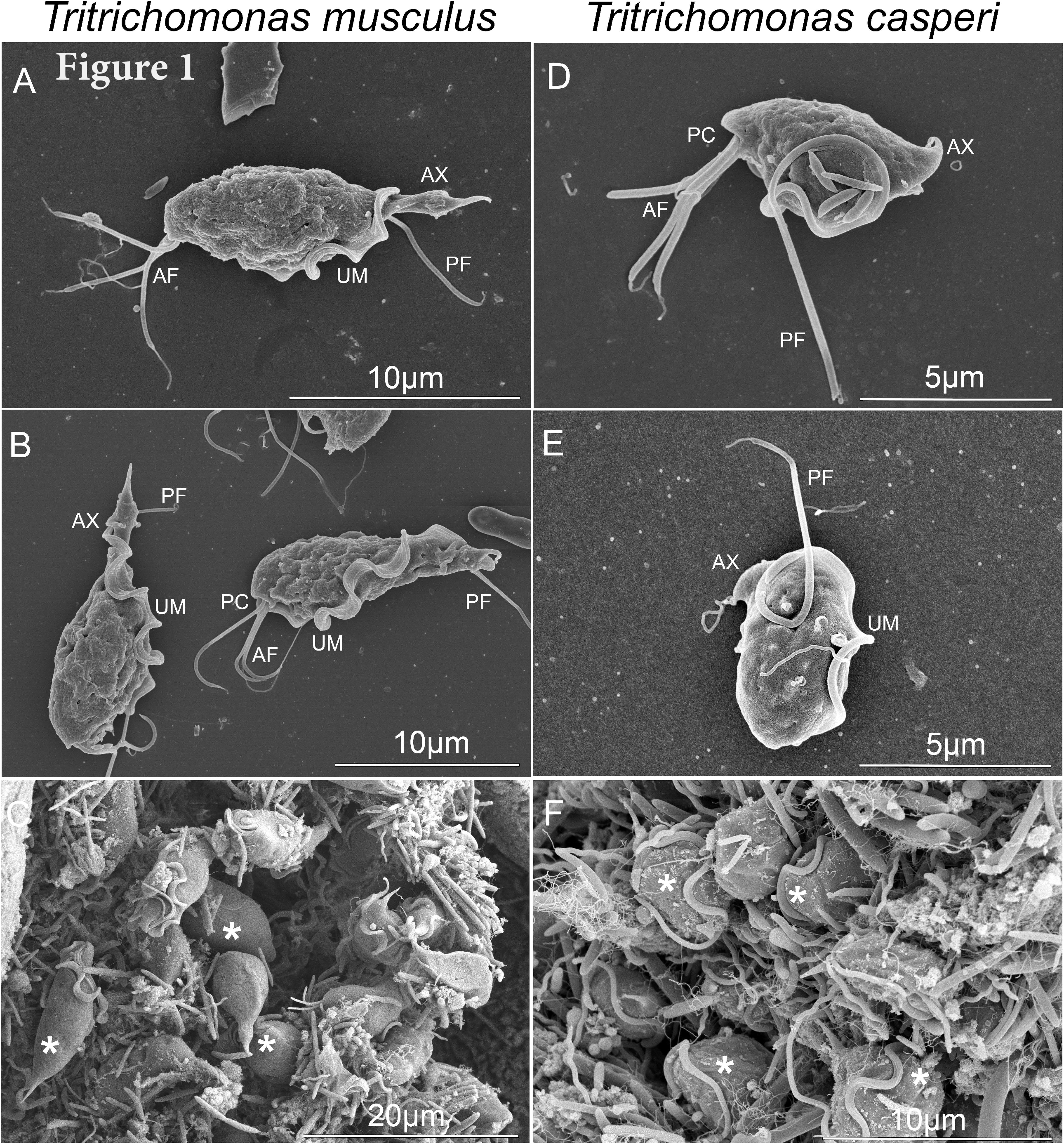
Scanning electron microscopy of *T. musculus* (A, B, and C) and *T. casperi* (D, E, and F), isolated from the cecum of laboratory mice (AF: anterior flagella, PC: periflagellar canal, UM: undulating membrane, AX: axostyle, and PF: posterior flagellum; *: protist). A, B, D, E are sorted cells; C, and F are *in situ* cecum preparations.

DNA was extracted from FACS-purified protists and two sets of PCR primers were applied to amplify a large fragment (1365 nucleotides) of the *18S rRNA* gene array, as well as the trans-*ITS* (267 nucleotides). For both protists, the two sets of pan-parabasalid primers generated electropherograms that were of high quality, with a single sequence type resolved, and no ambiguous bases identified. Phylogenetic analyses at the *18S rRNA* gene sequence confirmed that *T. casperi* was divergent from both *T. musculus*, and the highly similar sequence type annotated in GenBank as *T. rainier* (Acc No. MH370486) (Fig. 2A). A trichomonad isolated from laboratory mice in Russia, labeled with accession number MT804340.1, was 100% identical to *T. casperi* at the *18S rRNA* gene. *Tritrichomonas musculus* and *T. rainier* were distinct from each other and possessed two SNPs and one indel that differentiated the two protists at the SSU rRNA gene array within the V8 and V9 region (Fig. 2B). In addition, the more variable *ITS* marker provided additional support indicating that *T. rainier* likely represents a separate species (Fig. 3B). Studies are ongoing to identify mice infected with *T. rainier* to FACS-purify for ultrastructural characterization and species designation.

**FIGURE 2.**
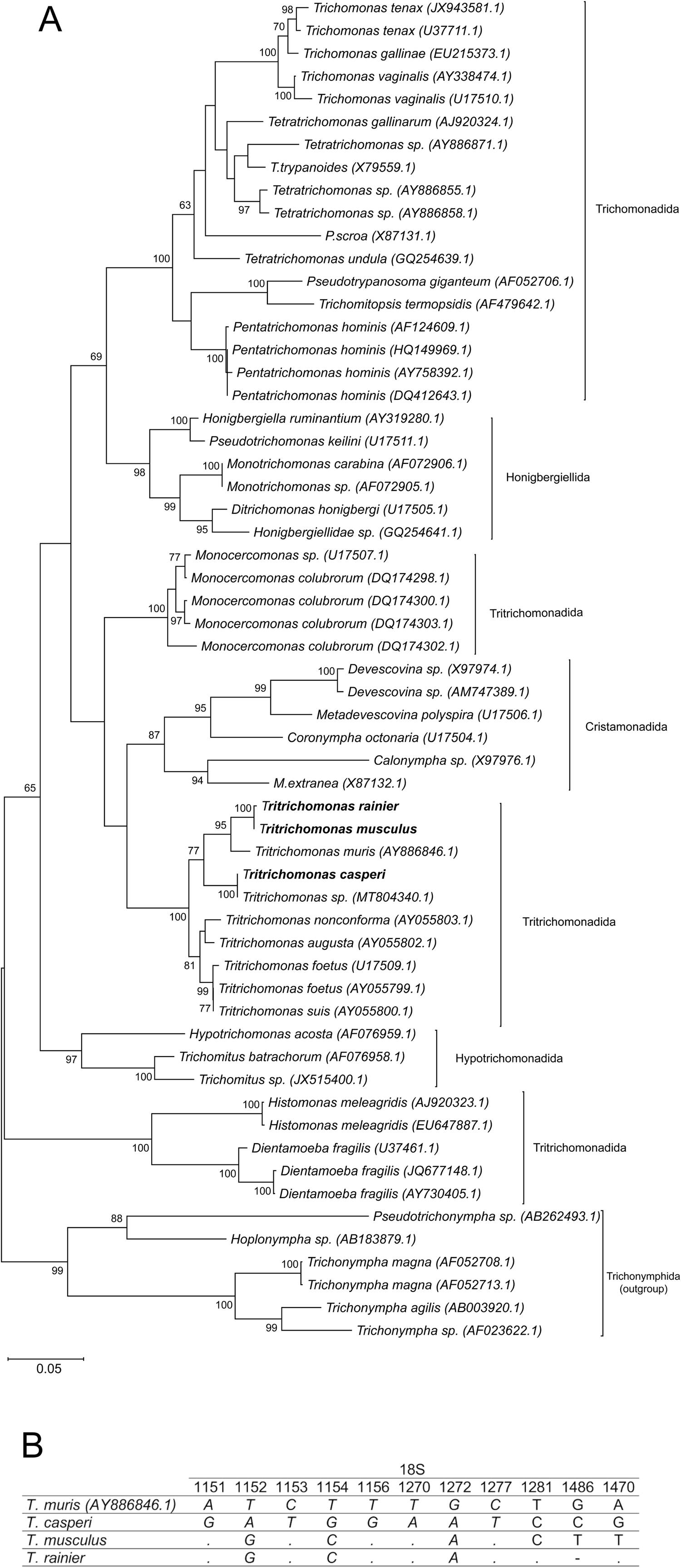
Characterization of trichomonad mouse commensals at the *18S rRNA* gene array. **A**. Phylogenetic tree showing *T. musculus, T. casperi*, and *T. rainier* relative to other trichomonad species. *18S rRNA* sequences deposited under Genbank Accession numbers ON907819, ON927245, and ON927247, respectively. DNA from purified *T. musculus, T. casperi*, and *T. rainier* was amplified at the *18S rRNA* gene array using primers PF1 and FAD4 (Keeling 2002). All sequences downloaded from NCBI are labelled with accession numbers. Sequences were aligned by MUSCLE using default settings in Geneious Prime v.2022.02. Phylogenetic tree was generated in MEGA X by Maximum Likelihood statistical method, 1000 bootstrap replications, and using the Tamura-Nei model. **B**. SNP sites of *T. musculus, T. casperi*, and *T. rainier* at the V8 through V9 region of the *18S rRNA* gene array that are distinct from *T. muris*. “.” indicates the SNP at the position identified is identical to that of *T. muris*, “-” denotes an INDEL.

**FIGURE 3.**
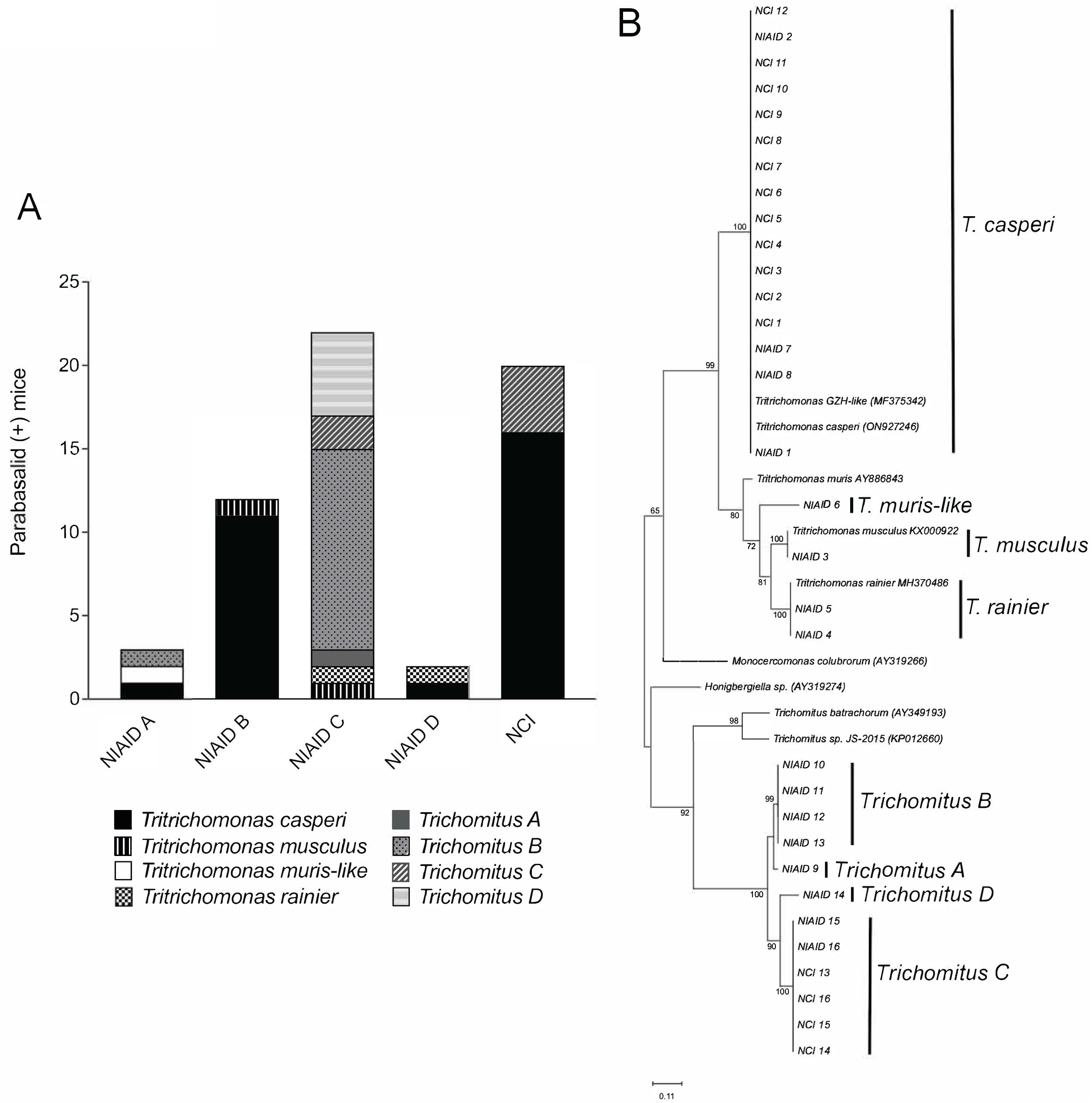
Diversity of parabasalids in laboratory mice across NIH based on trans-*ITS* sequencing. **A**. Distribution of 8 parabasalid sequence types across 5 different animal facilities. **B**. Phylogenetic tree with curated sequences from this study and sequences downloaded from NCBI labelled with accession numbers. Trans-*ITS* sequences were generated using pan-parabasalid primers anchored in the *18S* and *28S* rRNA gene array that amplify across the *ITS* (Chudnovskiy et al. 2016). Sequences were aligned using Clustal Omega (default settings) in Geneious Prime v.2022.02. The phylogenetic tree was generated in MEGA X by the Maximum Likelihood statistical method, 1000 bootstrap replications, using the Tamura-Nei model.

Overall, the tritrichomonas parasites of mice were monophyletic and formed a distinct clade relative to other trichomonads (Figure 2A). Their diversification into different species is arguably an example of sympatric speciation, but the factors that have led to this diversification remain enigmatic. Although these species were isolated from the cecum, their distribution throughout the GI tract was not uniform, and may indicate micro-niche adaptations to specific oxygen tensions, bacterial community structure, mucous, or various metabolites concentrated in separate ecological sites along the gastrointestinal tract that drive site specificity and speciation, as is known to occur, for example, among 8 *Eimeria* species of chickens (Barta et al. 1997). For trichomonads, another possibility driving such diversification is the acquisition of specific bacterial endosymbionts that provide metabolites, amino acids or nitrogen-derived compounds that confer an altered biological potential for micro-niche adaptation (Hongoh et al. 2008).

### Parabasalid Diversity Across NIH Animal Facilities

One animal facility within NCI and four animal facilities within NIAID (NIH, Bethesda, MD) were screened for the presence of parabasalids by fecal PCR DNA-Seq analyses of the PCR population using the trans-*ITS* locus. In total, 135 mouse stool samples collected from 135 separate cages across 8 rooms at the NIH were screened. The top BLAST results in each of the facilities sampled are displayed in Figure 3A, demonstrating that each facility had at least one parabasalid sequence type present. One animal facility on the NIH campus had the greatest diversity of parabasalid sequence types, with the dominant sequence type branching closely with *Trichomitus* (Acc ID AY349193 and KP01660). For a select number of the fecal PCR-positive mice, caecum was isolated in order to characterize the protists present by microscopy. *Tritrichomonas casperi* was the dominant sequence type recovered from mice in two different animal facilities, and it was present in four out of the five facilities screened. *Tritrichomonas musculus* and *T. rainier* were also detected across three facilities, whereas *T. muris* was not. Phylogenetic analysis at the trans-*ITS* gene array identified 8 distinct parabasalid sequence types (Fig. 3B). At least 4 sequence types resolved most closely with *Trichomitus* parasites, which are known to infect a variety of reptiles, including snakes (Dimasuay and Rivera 2013). The *Tritrichomonas* species formed a monophyletic clade of 4 distinct sequence types.

Although not the focus of this investigation, a number of mice were co-infected with two parabasalids. Thus, setting up molecular assays to screen for infection, as well as co-infection, will be important to identify the interactions that exist among these commensal trichomonads to ascertain how the presence of each individually, or collectively, impact the biochemical, immunological and physiological phenotypes that can be resolved during colonization with this diverse array of related protists.

The finding that *T. rainier-*derived succinate activates a type 2 immune response suggests that metabolites are a worthwhile avenue to investigate to determine how metabolites from these commensals impact immune cell development and/or cause shifts in bacterial community structure (Nadjsombati et al. 2018). Interestingly, one study showed that a high fiber diet facilitated the colonization of *T. musculus* compared to a high fat diet, corroborating the importance of metabolites involved in fiber digestion. The same study also highlighted how the administration of a broad-spectrum antibiotic in mice facilitated *T. musculus* colonization, regardless of diet, suggesting that bacteria in the gut ecosystem are an important consideration in the successful establishment of trichomonad infections in the gut (Wei et al. 2020). Substantial research on protist-bacteria symbioses in the termite gut provides an interesting perspective on how protists and bacteria have co-evolved to be essential components of nutrient acquisition in termites (Peterson, Stewart, and Scharf 2015). Whether this phenomenon is occurring in the intestinal lumen of mice is not known, and likely serves as an important factor driving sympatric speciation for this order of single-celled protists that naturally colonize rodents.

## CONCLUSION

A total of 8 distinct parabasalid sequence types were identified, highlighting an extant genetic diversity of protists infecting laboratory mice across the NIH. *Tritrichomonas casperi* was the dominant parabasalid identified, but *Tritrichomonas rainier* and *Tritrichomonas musculus* were detected in two and one mice sampled, respectively. Phylogenetic analyses at the *18S SSU rRNA* and trans-*ITS* gene array support the conclusion that these protists exist as distinct species. The morphological and ultrastructural data further substantiates the conclusion that *T. casperi* and *T. musculus* are separate species, distinct from *T. muris*. Moreover, *T. muris* was not detected in any of the 135 mouse fecal samples screened, suggesting that it is not the primary flagellate colonizing laboratory mice, as suggested previously (Escalante et al. 2016; Howitt et al. 2016). Although one sample shared 90% sequence similarity with *T. muris* at the trans-*ITS* region, it branched more closely with *T. musculus* than with *T. muris* (Fig 3B) providing additional evidence that other mouse commensals within this genus remain to be described.

## TAXONOMIC SUMMARY

**Taxonomic Summary:** Eukaryota; Metamonada; Parabasalia; Tritrichomonadida; Tritrichomonadidae; *Tritrichomonas musculus*

**Type Host:** *Mus musculus*

**Other hosts:** Unknown

**Type Habitat:** Cecum of rodent

**Type locality:** NIH Animal Facility, Bethesda, MD, USA

**Etymology:** Named after the host genus

**Gene sequence:** The DNA sequences are deposited in Genbank. Accession numbers are ON907819/KX000921 (*18S rRNA*) and KX000921 (trans-*ITS*).

**Zoobank registration:** 857778FE-4EE2-4242-A0B3-F92492130E1E

**Type Material:** Specimens are deposited at the Department of Invertebrate Zoology Collections, National Museum of Natural History, Smithsonian Institution, Washington, DC in the form of SEM stubs and TEM resin blocks containing FACS purified *T. musculus* under the USNM code (XXX).

**Description of *T. musculus*, n. sp. (Fig. 1A, 1B):** A multi-flagellated commensal protist 10 μm in size was isolated from the cecum of laboratory mouse (*Mus musculus*). It had 3 anterior and 1 posterior flagella and possessed an undulating membrane that protruded ~1 μm from the surface of the protist and spiraled in a ribbon-like manner from the periflagellar canal to the axostyle. In liquid medium, *T. musculus* moved in a smooth, gliding fashion with the undulating membrane beating in a rhythmic manner. The *18S rRNA* and trans-*ITS* sequences were distinct from *T. muris*.

**Taxonomic Summary:** Eukaryota; Metamonada; Parabasalia; Tritrichomonadida; Tritrichomonadidae; *Tritrichomonas; casperi*

**Type Host:** *Mus musculus*

**Other hosts:** Unknown

**Type Habitat:** Cecum of rodent

**Type locality:** NIH Animal Facility, Bethesda, MD, USA

**Etymology:** *T. casperi* was discovered independently by flow cytometry in the Grigg lab at the NIH and in the Howitt Lab at Stanford University. During discussions between the two labs, it was remarked that *T. casperi* often existed as a smaller “shadow” when purifying *T. musculus* from co-infected mice. Hence, it was considered a “ghost” and named after “Casper the ghost”

**Gene sequence:** The DNA sequences were deposited in Genbank. Accession numbers are ON927245 (*18S rRNA*) and ON927246 (trans-*ITS*).

**Zoobank registration:** 6EF2A037-7429-4B93-A236-065B95FBE4E8

**Type Material:** Specimens are deposited at the Department of Invertebrate Zoology Collections, National Museum of Natural History, Smithsonian Institution, Washington, DC in the form of SEM stubs and TEM resin blocks containing FACS purified *T. casperi* under the USNM code (XXX).

**Description of *T. casperi*, n. sp. (Fig. 1C, 1D):** A multi-flagellated commensal protist 5 μm in size was isolated from the cecum of a laboratory mouse (*Mus musculus*). It had 3 anterior and 1 posterior flagella and a small undulating membrane that protruded 0.3 μm from the surface of the protist and ran along the dorsal flank in a more linear fashion than that observed for *T. musculus* before pivoting away from the main ovaloid body to form the posterior flagellum. In liquid medium, *T. casperi* moved in a stochastic, apparently erratic fashion that was distinct from *T. musculus*. The *18S rRNA* and trans-*ITS* sequences were likewise distinct from both *T. musculus* and *T. muris*.

## ACKNOWLEDGEMENTS

This work was supported by the Intramural Research Program of the National Institute of Allergy and Infectious Diseases (NIAID) at the National Institutes of Health. The authors would like to thank Andrea Paun and members of the Grigg lab and the NIH community for providing access to mouse fecal samples from the five animal facilities studied. Dr. Miriam Merad, Dr. Yasmine Belkaid, Dr. Jakob von Moltke and Dr. Michael Howitt for ongoing discussions on trichomonads infecting mice, and Dr. Jakob von Moltke for DNA from *Tritrichomonas rainier*.

## Ethics statement

All rodent work was approved under the animal study protocol LPD 22E, reviewed and approved by the Animal Care and Use Committee of the Intramural Research Program of the National Institute of Allergy and Infectious Diseases, National Institutes of Health. All datasets generated for this study are available upon request to the corresponding author.

## REFERENCES

Bar, A. K., N. Phukan, J. Pinheiro, and A. Simoes-Barbosa. 2015. “The Interplay of Host Microbiota and Parasitic Protozoans at Mucosal Interfaces: Implications for the Outcomes of Infections and Diseases.” PLoS Negl Trop Dis 9 (12): e0004176. https://doi.org/10.1371/journal.pntd.0004176. https://www.ncbi.nlm.nih.gov/pubmed/26658061.

Barta, J. R., D. S. Martin, P. A. Liberator, M. Dashkevicz, J. W. Anderson, S. D. Feighner, A. Elbrecht, A. PerkinsBarrow, M. C. Jenkins, H. D. Danforth, M. D. Ruff, and H. ProfousJuchelka. 1997. “Phylogenetic relationships among eight Eimeria species infecting domestic fowl inferred using complete small subunit ribosomal DNA sequences.” Journal of Parasitology 83 (2): 262–271. https://doi.org/10.2307/3284453. https://www.ncbi.nlm.nih.gov/pubmed/9105308.

Cepicka, I., Hampl, V., and Kulda, J. 2010. “Critical taxonomic revision of parabasalids with description of one new genus and three new species.” Protist 161:400–433. https://doi.org/10.1016/j.protis.2009.11.005. https://www.ncbi.nlm.nih.gov/pubmed/20093080.

Ceza, V., T. Panek, P. Smejkalova, and I. Cepicka. 2015. “Molecular and morphological diversity of the genus Hypotrichomonas (Parabasalia: Hypotrichomonadida), with descriptions of six new species.” Eur J Protistol 51 (2): 158–72. https://doi.org/10.1016/j.ejop.2015.02.003. https://www.ncbi.nlm.nih.gov/pubmed/25855142.

Chudnovskiy, A., A. Mortha, V. Kana, A. Kennard, J. D. Ramirez, A. Rahman, R. Remark, I. Mogno, R. Ng, S. Gnjatic, E. D. Amir, A. Solovyov, B. Greenbaum, J. Clemente, J. Faith, Y. Belkaid, M. E. Grigg, and M. Merad. 2016. “Host-Protozoan Interactions Protect from Mucosal Infections through Activation of the Inflammasome.” Cell 167 (2): 444–456 e14. https://doi.org/10.1016/j.cell.2016.08.076. https://www.ncbi.nlm.nih.gov/pubmed/27716507.

Edgar, R. C. 2004. “MUSCLE: multiple sequence alignment with high accuracy and high throughput.” Nucleic Acids Res 32 (5): 1792–7. https://doi.org/10.1093/nar/gkh340. https://www.ncbi.nlm.nih.gov/pubmed/15034147.

Escalante, N. K., P. Lemire, M. Cruz Tleugabulova, D. Prescott, A. Mortha, C. J. Streutker, S. E. Girardin, D. J. Philpott, and T. Mallevaey. 2016. “The common mouse protozoa Tritrichomonas muris alters mucosal T cell homeostasis and colitis susceptibility.” J Exp Med 213 (13): 2841–2850. https://doi.org/10.1084/jem.20161776. https://www.ncbi.nlm.nih.gov/pubmed/27836928.

Hongoh, Y., V. K. Sharma, T. Prakash, S. Noda, H. Toh, T. D. Taylor, T. Kudo, Y. Sakaki, A. Toyoda, M. Hattori, and M. Ohkuma. 2008. “Genome of an Endosymbiont Coupling N-2 Fixation to Cellulolysis Within Protist Cells in Termite Gut.” Science 322 (5904): 1108–1109. https://doi.org/10.1126/science1165578. <Go to ISI>://WOS:000260867700039.

Howitt, M. R., S. Lavoie, M. Michaud, A. M. Blum, S. V. Tran, J. V. Weinstock, C. A. Gallini, K. Redding, R. F. Margolskee, L. C. Osborne, D. Artis, and W. S. Garrett. 2016. “Tuft cells, taste-chemosensory cells, orchestrate parasite type 2 immunity in the gut.” Science 351 (6279): 1329–33. https://doi.org/10.1126/science.aaf1648. https://www.ncbi.nlm.nih.gov/pubmed/26847546.

Jackson, K. M., C. Schwartz, J. Wachter, P. A. Rosa, and P. E. Stewart. 2018. “A widely conserved bacterial cytoskeletal component influences unique helical shape and motility of the spirochete Leptospira biflexa.” Mol Microbiol 108 (1): 77–89. https://doi.org/10.1111/mmi.13917. https://www.ncbi.nlm.nih.gov/pubmed/29363884.

Keeling, P. J. 2002. “Molecular phylogenetic position of Trichomitopsis termopsidis (Parabasalia) and evidence for the Trichomitopsiinae.” European Journal of Protistology 38 (3): 279–286. https://doi.org/Doi10.1078/0932-4739-00874. <Go to ISI>://WOS:000179057800007.

Lukes, J., C. R. Stensvold, K. Jirku-Pomajbikova, and L. Wegener Parfrey. 2015. “Are Human Intestinal Eukaryotes Beneficial or Commensals?” PLoS Pathog 11 (8): e1005039. https://doi.org/10.1371/journal.ppat.1005039. https://www.ncbi.nlm.nih.gov/pubmed/26270819.

Nadjsombati, M. S., J. W. McGinty, M. R. Lyons-Cohen, J. B. Jaffe, L. DiPeso, C. Schneider, C. N. Miller, J. L. Pollack, G. A. Nagana Gowda, M. F. Fontana, D. J. Erle, M. S. Anderson, R. M. Locksley, D. Raftery, and J. von Moltke. 2018. “Detection of Succinate by Intestinal Tuft Cells Triggers a Type 2 Innate Immune Circuit.” Immunity 49 (1): 33–41 e7. https://doi.org/10.1016/j.immuni.2018.06.016. https://www.ncbi.nlm.nih.gov/pubmed/30021144.

Nieves-Ramirez, M. E., O. Partida-Rodriguez, I. Laforest-Lapointe, L. A. Reynolds, E. M. Brown, A. Valdez-Salazar, P. Moran-Silva, L. Rojas-Velazquez, E. Morien, L. W. Parfrey, M. Jin, J. Walter, J. Torres, M. C. Arrieta, C. Ximenez-Garcia, and B. B. Finlay. 2018. “Asymptomatic Intestinal Colonization with Protist Blastocystis Is Strongly Associated with Distinct Microbiome Ecological Patterns.” mSystems 3 (3). https://doi.org/10.1128/mSystems.00007-18. https://www.ncbi.nlm.nih.gov/pubmed/29963639.

Peterson, B. F., H. L. Stewart, and M. E. Scharf. 2015. “Quantification of symbiotic contributions to lower termite lignocellulose digestion using antimicrobial treatments.” Insect Biochem Mol Biol 59: 80–8. https://doi.org/10.1016/j.ibmb.2015.02.009. https://www.ncbi.nlm.nih.gov/pubmed/25724277.

Sievers F, Barton GJ, Higgins DG. 2020. “Mutliple Sequence Alignment.” In Bioinformatics, edited by GD Bader AD Baxevanis, DS Wishart, 227–250.

Sievers, F., and D. G. Higgins. 2018. “Clustal Omega for making accurate alignments of many protein sequences.” Protein Sci 27 (1): 135–145. https://doi.org/10.1002/pro.3290. https://www.ncbi.nlm.nih.gov/pubmed/28884485.

Sievers, F., A. Wilm, D. Dineen, T. J. Gibson, K. Karplus, W. Li, R. Lopez, H. McWilliam, M. Remmert, J. Soding, J. D. Thompson, and D. G. Higgins. 2011. “Fast, scalable generation of highquality protein multiple sequence alignments using Clustal Omega.” Mol Syst Biol 7: 539. https://doi.org/10.1038/msb.2011.75. https://www.ncbi.nlm.nih.gov/pubmed/21988835.

Solaymani-Mohammadi, S. 2022. “Mucosal Defense Against Giardia at the Intestinal Epithelial Cell Interface.” Front Immunol 13: 817468. https://doi.org/10.3389/fimmu.2022.817468. https://www.ncbi.nlm.nih.gov/pubmed/35250996.

Stecher, G., K. Tamura, and S. Kumar. 2020. “Molecular Evolutionary Genetics Analysis (MEGA) for macOS.” Mol Biol Evol 37 (4): 1237–1239. https://doi.org/10.1093/molbev/msz312. https://www.ncbi.nlm.nih.gov/pubmed/31904846.

Treuting, P. M., C. B. Clifford, R. S. Sellers, and C. F. Brayton. 2012. “Of mice and microflora: considerations for genetically engineered mice.” Vet Pathol 49 (1): 44–63. https://doi.org/10.1177/0300985811431446. https://www.ncbi.nlm.nih.gov/pubmed/22173977.

Wei, Y., J. Gao, Y. Kou, L. Meng, X. Zheng, M. Liang, H. Sun, Z. Liu, and Y. Wang. 2020. “Commensal Bacteria Impact a Protozoan’s Integration into the Murine Gut Microbiota in a Dietary Nutrient-Dependent Manner.” Appl Environ Microbiol 86 (11). https://doi.org/10.1128/AEM.00303-20. https://www.ncbi.nlm.nih.gov/pubmed/32198171.

